# Sex chromosomes and sex hormones contribute jointly and independently to sex biases in cardiac development

**DOI:** 10.64898/2026.01.09.696197

**Authors:** Daniel F. Deegan, Gennaro Calendo, Priya Nigam, Raza Naqvi, Arthur P. Arnold, Nora Engel

## Abstract

Congenital heart defects are highly sex-biased, and although the regulatory networks underlying the differentiation of cardiac progenitors are well established, the sex differences during early stages of cardiogenesis have been largely understudied. Traditionally, sex differences are attributed to the gonadal sex hormones, but the unequal sex chromosome complement (i.e., XX *versus* XY) contributes to sex-biased gene expression from soon after fertilization and across the lifespan. We leveraged the Four Core Genotypes (FCG) mouse model to investigate gonadal and sex chromosome effects on the transcriptome during cardiac development. We found significant sex-biased gene expression at all developmental stages, including in 10.5 dpc (days post-coitum) embryos, before the gonads have formed. Our studies reveal that sex differences in the transcriptome are governed by both sex chromosome and gonadal sex hormone effects and their interactions in 16.5 dpc, neonates, and adults, to differing degrees depending on the stage. Transcriptional and epigenetic factors were among the differentially expressed genes, suggesting the existence of sex-specific subnetworks. The adult cardiac epigenome was also investigated for differential H3K27Ac enrichment, revealing sex-biased chromatin states containing transcription factor binding sites regulated by sex chromosomes, gonadal hormones, and their combined effects. Several of these differentially enriched regions were found to overlap with known cardiac enhancers and correlated with differential expression of typical cardiac-specific genes.

## Introduction

After years of assuming that gonadal hormones were the sole contributors to sex differences, we now know that the sex chromosome complement independently imposes biases on gene expression and the epigenome beginning in male and female embryos soon after fertilization. This revelation may provide clues to why certain congenital heart defects are highly sex-biased^1,2^. Congenital heart defects are associated with disruptions in the differentiation of cardiac progenitors, so major efforts have been aimed at elucidating the regulatory networks underlying cardiogenesis^3-7^. However, sex differences have been mostly overlooked in developmental processes. Identifying the molecular basis for sex differences throughout heart development can provide insight into sex biases in congenital heart defects. It also has the potential to yield information on sex differences in cardiovascular health and disease across the lifespan, including both susceptibility and protective factors that can be therapeutic targets.

We and others have shown that sex-biased gene expression is evident soon after fertilization in response to the sex chromosome complement^8-11^. In mouse embryonic stem cells, some of these sex biases equalize or disappear after differentiation, but new ones emerge, including differences in transcriptional and epigenetic factors^12^. This implies that after gonad formation, sex hormones act on a cardiac transcriptome and an epigenome that are already distinct between the sexes. In addition, sex chromosome-linked genes continue to have an independent effect on gene expression^13^, either directly or via downstream autosomal effectors of sex chromosome-encoded regulatory factors. We have previously shown that sex-biased gene expression exists in early cardiac development before and after gonad formation^14^, and proteomic studies have confirmed that these differences are present at the protein level as well^15^. In fact, some expression biases found at early stages of cardiogenesis are conserved in adult cardiomyocytes^14^. This suggests that there are mechanisms that maintain the sexual identity imprinted upon the genome by sex chromosome-linked genes during early development in addition to those programmed at later stages.

After gonad formation, ovarian and testicular hormones begin to circulate and contribute to organogenesis, leading to permanent effects on the transcriptome and epigenome. In addition, sex hormones have cyclical and reversible effects after puberty. In the heart, both estrogens and androgens and their receptors are essential for normal cardiac function. For example, estrogens promote the survival of cardiac myocytes^16^. Similarly, endogenous testosterone aids in preventing cardiomyocyte apoptosis and cardiac fibrosis, resulting in a strong overall protective effect on the heart^17^. The effects of the sex hormones, however, change throughout development and can differ greatly between males and females. The risk of cardiovascular disease (CVD) in women is strongly correlated with menopause, with risk rising significantly post-menopause. This can be explained at least in part by the reduction of endogenous estrogens^18^, but does not completely account for the fact that hormone replacement therapy does not seem to restore protection^16^. Interestingly, the effect of estrogens on CVD in men is still inconclusive. Low levels of testosterone are associated with a higher risk of cardiovascular disease in older men compared to their younger counterparts^18^. The impact of testosterone on cardiovascular disease in women, however, is unclear. Even with these lingering questions, the influence of sex hormones on cardiac function and health is unequivocal.

In addition to contributing independently to sex differences, sex chromosomes and sex hormones can also synergize or antagonize each other’s effects. However, because the two factors co-vary, unravelling their independent effects is challenging. Our hypothesis is that sex chromosomes and sex hormones have joint and independent roles in sex-biased gene expression and epigenetics throughout cardiac development, with consequences for the sex differences observed in adults at molecular and functional levels.

To systematically evaluate the relative importance, effect size and dynamics of each factor at the transcriptional level, we leveraged the Four Core Genotypes (FCG) mouse model^19^, in which the gonad type is independent of the sex chromosome complement. We quantified RNA expression across cardiac development in the FCG model, beginning in embryos at 10.5 days post-coitum (dpc), before gonad formation. We find a dynamic pattern of increasing sex-biased expression, ranging from few genes at early embryonic stages, to hundreds of genes at later stages. In addition, we identify subsets of differentially expressed genes and sex-specific chromatin modifications that are solely dependent on the sex chromosome constitution or the gonadal type, as well as those that depend on the interaction of both factors.

## Methods

### Animals

Mouse studies were performed under approval of the Temple University Institutional Animal Care and Use Committee. Mice were maintained under standard conditions of 12-h light and dark cycles (22°C ± 1°C, with food and water *ad libitum*). We used the Four Core Genotypes (FCG) mouse model^19,20^ on a C57BL/6J B6 background (B6.Cg-Tg(Sry)2Ei *Sry*^*dl1Rlb*^ T(XTmsb4x-Hccs;Y)1Dto/ArnoJ, Jackson Laboratories stock 010905; backcross generation greater than 20), derived from the Arnold lab at UCLA. The founder males in this model have the *Sry* gene deleted from the Y chromosome and inserted as a transgene on chromosome 3 (XY^-^*Sry*^+^, referred to hereafter as XYM). Crossing the XYM with a wild-type XX female results in XX and XY mice with ovaries (designated XXF and XYF) and XX and XY mice with testes (designated XXM and XYM), dependent solely on whether the *Sry* transgene is inherited and independently of the presence of the Y chromosome. Thus, the gonadal type of the progeny (testes or ovaries) is independent of the sex chromosome constitution. Analyzing these four core genotypes allows distinguishing between sex chromosome and sex hormone dependent effects (**Figure 1**).

**Figure 1.**
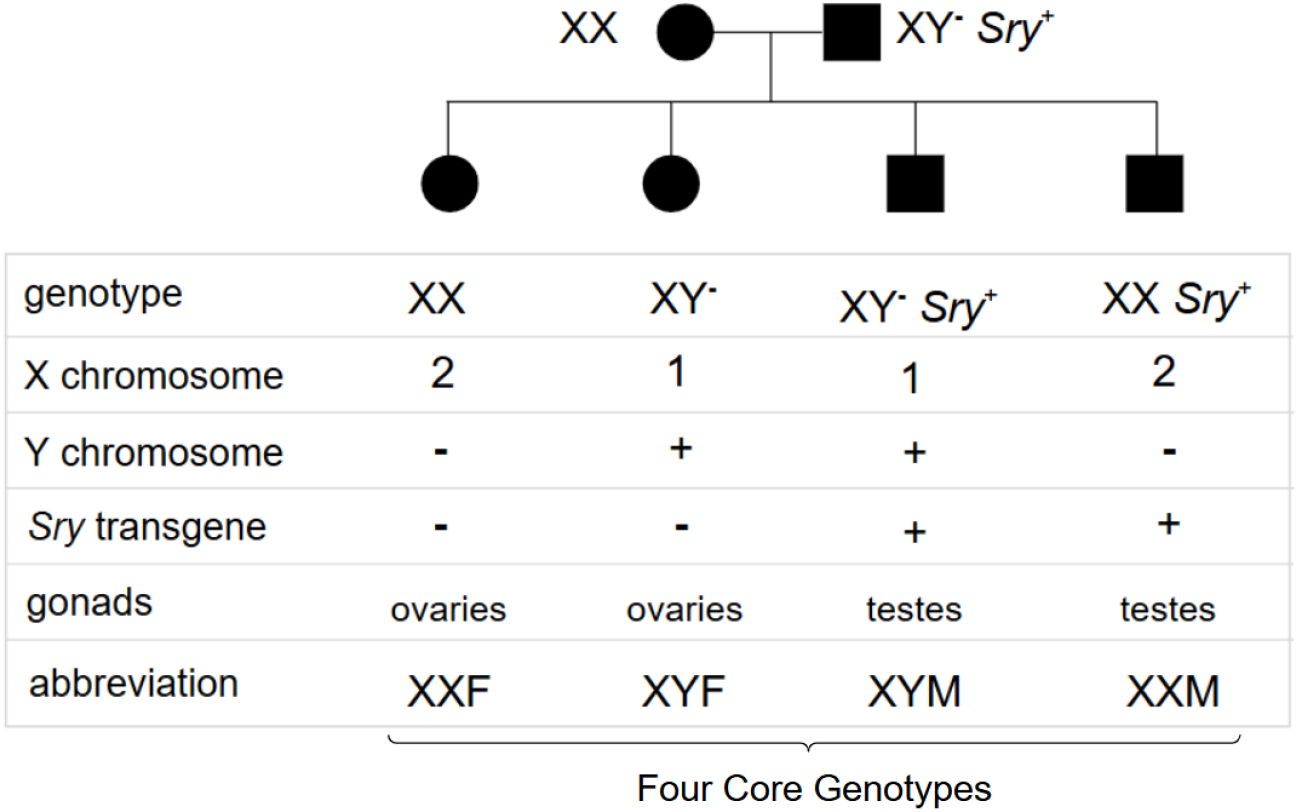
The Four Core Genotypes (FCG) mouse model.

The FCG model used here involves a Y chromosome onto which 9 X-linked genes were translocated, during backcross of the model from strain MF1 to C57BL/6J^21^. To avoid any biases introduced from the potential increased expression of these translocated genes, differential expression of these genes was excluded from all analyses. Numerous XX-XY differences discovered using FCG mice have, however, been found to be caused by higher expression of X genes in females or Y genes in males (*not* the 9 translocated genes), reinforcing the idea that most XX-XY differences in FCG mice reflect typical XX-XY differences in wild-type mice^22,23^.

The founder XY^-^*Sry*^+^ males (XYM) on a C57BL/6J background were used for timed matings with C57BL/6J females, with the detection of a vaginal plug counted as day 0.5. Hearts were obtained from the four genotypes (XXM, XXF, XYM, XYF) at four different stages: 10.5 dpc, 16.5 dpc, neonates and adults. Hearts were dissected and snap-frozen in liquid nitrogen and then stored at -80°C for RNA, DNA, and chromatin extraction.

### Genotyping

DNA was extracted from embryos or tails of neonatal and adult mice and genotype was determined by PCR using primers for *Sry* and *Ymt*, with *Kcnq1* as a positive control. PCR products were then run on a 2% agarose gel. Gonadal males exhibit a positive result for *Sry*, and positive or negative for *Ymt* reveals the presence or absence of a Y chromosome^20^. PCRs were performed using DreamTaq Green PCR Mastermix (2x) (Thermofisher). Primer sequence, annealing temperature, and product size are listed in **Supplemental Table 1**.

### RNA isolation and quality control

RNA was extracted from tissues using the Quick RNA Microprep Kit (Zymo) for hearts from 10.5 dpc embryos and Quick RNA Miniprep Kit (Zymo) for 16.5 dpc and neonatal hearts. For adult hearts, RNA was extracted with the Monarch Total RNA Miniprep Kit (New England Biolabs). The RNA was treated with DNase using the TURBO DNA-free Kit (Thermofisher). RNA integrity was analyzed using the Agilent BioAnalyzer 2100 and Agilent’s RNA 6000 Nano Kit. Only samples with an RNA Integrity Number (RIN) score greater than 7.0 were considered high integrity and moved on to sequencing.

### RNA-sequencing and data analysis

A total of three biological replicates for each genotype and stage were obtained for RNA-seq (48 samples). Samples were sequenced in two batches. At the Fox Chase Cancer Center sequencing core, libraries were prepared using TakaRa Bio’s SMARTer pico-input kit (Cat# 634940) for low-input RNA samples. Sequencing was performed using the following parameters: 1X single reads with a 1x75bp read length at a read depth of 30 million reads per sample. All samples were sequenced using an Illumina NextSeq 500 sequencer. The second batch was sequenced at Novogene (150 pair-end). To harmonize the sequencing, only the first read of the paired-end sequencing was used and was hard-clipped to 80bp using trim_galore^24^. Otherwise, samples were processed uniformly. RNA-seq datasets were aligned to mm10 with STAR version 2.7.0^25^. Gene counts were generated using featureCounts^26^ with the inbuilt mm10 annotation. Gene count matrices were imported into edgeR^27^ for downstream differential expression analysis. Read counts were filtered to remove lowly expressed genes using the ‘filterByExpr()’ function in edgeR. Trimmed mean of the M-value (TMM) normalization factors were computed for each library. A linear model was fit with the glmQLFit() function to determine sex-chromosome, sex-hormone, and developmental stage independent and interaction effects, controlling for batch. Additionally, a stage-stratified analysis was performed to determine independent and interaction effects of sex chromosomes and gonadal hormones within developmental stage, controlling for batch. Differential expression was tested for each coefficient using the glmQLFTest() function. Genes with FDR<0.05 were considered significantly differentially expressed (**Supplemental Table 2**,**3**,**4; Supplemental Figure 1**).

Gene ontology (GO) analysis was performed using the up/down-regulated genes in each contrast. Gene set testing was performed against the HALLMARK gene sets from MSigDB using a competitive gene set test, ‘camera()’, implemented in the ‘edgeR’ package (**Supplemental Tables 5**,**6**).

### CUT&RUN

Two biological replicates of adult heart samples from each of the four genotypes were processed with the CUT&RUN Assay Kit (Cell Signaling Technologies®). Approximately 5mg of heart tissue was minced into small pieces using a scalpel, fixed in 0.1% formaldehyde for 5 minutes and homogenized. Samples were prepared using antibody against Acetyl-Histone H3 (Lys27) (Cell Signaling Technologies®). A 1:10 dilution of this antibody (10µl) was used per each sample. Positive and negative control antibodies were included in each experiment. Samples were incubated at 4°C overnight on a nutator. The input sample was processed according to the protocol and sonicated using the Covaris® M220 Focused Ultrasonicator. The parameters for sonication were 20°C, 50 Peak Incident Power (PIP), 20% Duty Factor, 200 Cycles Per Burst, and 65 seconds. All samples, including the input, were purified using the DNA Purification Buffers and Spin Columns (ChIP, CUT&RUN) kit (Cell Signaling Technologies®).

Quantitative PCR (qPCR) was used to assess the percent input of each sample to reveal if the enrichment was successful. The protocol outlined in the CUT&RUN Assay Kit was followed accordingly with the following modifications: PowerUp™ SYBR Green Master Mix (Catalog #A25742) from Thermofisher was utilized and a 5-point serial dilution was used to calculate the standard curve (undiluted, 1:5, 1:25, 1:125, and 1:625). SimpleChIP® Mouse RPL30 Intron 2 Primers (Cell Signaling Technologies® Catalog #7015) were used to assess percent input. An experiment was considered successful if the Tri-Methyl Histone H3 (Lys4) (Catalog #9751) positive control percent input was > 1.0% and significantly greater than the percent input for the IgG Isotype (Catalog #66362) negative control.

CUT&RUN samples were sent to Novogene Corporation Inc. for sequencing. Library preparation was performed using KAPA Hyper Prep kit (Roche) per manufacturer’s recommendations. Briefly, adapters were ligated to DNA fragments that were then amplified with minimal PCR cycles. Library quantity and quality were assessed with Qubit 2.0 DNA HS Assay (ThermoFisher), Tapestation High Sensitivity D1000 Assay (Agilent Technologies), and QuantStudio ® 5 System (Applied Biosystems). Illumina® 8-nt dual-indices were used. Equimolar pooling of libraries was performed based on QC values and sequenced on an Illumina NovaSeq 6000 with a read length configuration of 150 PE for 80M PE reads (40M in each direction).

### CUT&RUN data processing

CUT&RUN experiments were performed on two biological replicates of adult hearts per genotype. Adapter trimming and quality control of raw paired-end reads was performed using fastp^28^. Trimmed reads were aligned to GRCm39 using Bowtie2^29^ in -- end-to-end mode with the following arguments, --no-mixed, --no-discordant, --dovetail, --very-sensitive. The resulting BAM files were filtered for properly paired alignments, reads mapping to mitochondrial chromosomes were removed, and duplicates were marked using samtools^30^. Filtered alignment files were then converted to bedgraph files using bedtools^31^. Peak calling was performed on bedgraphs using SEACR^32^ in ‘stringent’ mode. A consensus peak set was defined by taking the union of all peaks detected across groups, where the intersection of all peaks between replicate conditions per group defined groupwise peaks. featureCounts^26^ was used to generate an abundance count matrix of the consensus peak set. Differential abundance analysis was performed using edgeR to determine sex-chromosome independent, sex-hormone independent, and chromosome-hormone interaction effects. Peaks with FDR<0.05 were considered significantly differentially abundant.

### Differential peak abundance analysis

Peaks used for differential analysis were determined by taking the intersection of peak calls for each sample within each group. ‘featureCounts()’ was used to count reads from the filtered BAM files in the combined peak set. Read counts were then used in differential abundance testing after removing peaks with low counts across all samples (aveLog2CPM < 3). Differential abundance between conditions was assessed using the ‘edgeR::glmQLFTest()’. The design used in this analysis was “∼0 + group”, where each group represented one of the FCGs. This design facilitates comparisons between all combinations of groups. The contrasts assessed for differential abundance were: XXF *versus* XYF, XXM *versus* XYM, XYF *versus* XYM, XXF *versus* XXM, female *versus* male (average difference holding sex chromosomes constant) and XX *versus* XY (average difference holding hormonal status constant). Peaks were then annotated using ‘ChIPseeker::annotatePeak()’ against the GENCODE vM27 reference GTF; the default promoter region used in the ‘annotatePeak()’ function is –[3000, 3000] of the TSS (**Supplemental Table 7**).

### Motif analysis for differentially enriched H3K27Ac peaks

For each set of comparisons, peaks that were significantly up- or down-regulated (FDR<0.05) were extracted and motif analysis was performed with the XSTREME analysis implemented in the ‘MEME’ suite^33^. De novo motifs were compared to known motifs in the HOCOMOCO v11 core MOUSE database (**Supplemental Table 8**).

### Concordance analysis between RNA-seq and the CUT&RUN data

Prior to performing concordance analysis, CUT&RUN peak coordinates were lifted from GRCm39 to mm10.

Differentially expressed genes were correlated with their nearest upstream CUT&RUN peak from the CUT&RUN analysis. Upstream peaks were computed only for the differential expression analysis data from the stratified analysis on Adults for XX *versus* XY and Female *versus* Male (**Supplemental Table 9**).

### Overlap analysis of differentially abundant peaks with cardiac enhancers

A list of known cardiac enhancers was downloaded from SCREEN^34^ using the following annotation: “C57BL/6 heart tissue male adult 8 weeks”. Then, regions representing significantly differentially abundant peaks that are hormone-dependent (F *versus* M) or sex chromosome-dependent (XX *versus* XY) that also overlap a known cardiac enhancer were determined (**Supplemental Table 10**).

## Results

### Sex-biased gene expression is apparent throughout cardiogenesis

To systematically evaluate the relative contributions of sex chromosomes and sex hormones on gene expression during cardiac development, we used the Four Core Genotypes (FCG) mouse model in which the gonad type is independent of the sex chromosome complement (**Methods, Figure 1)**. We obtained hearts from mice of each of the four genotypes (XXF, XXM, XYF, XYM) at 10.5 and 16.5 dpc embryos, neonates, and adults and conducted RNA-sequencing for transcriptome analysis (**Figure 2**). Sex chromosome effects (SCEs) were determined by comparing XX and XY mice, whereas sex hormone effects (SHEs) were evaluated by comparing gonadal males to gonadal females.

**Figure 2.**
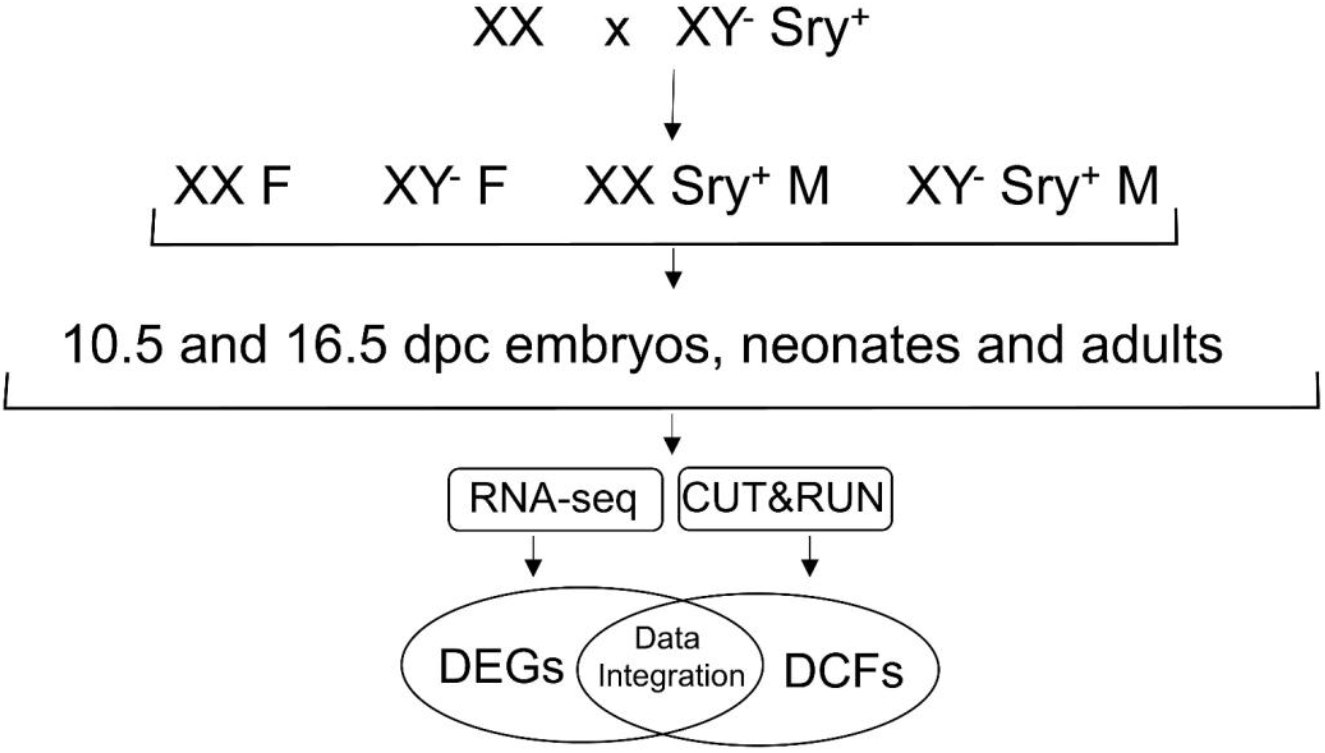
Experimental design. The Four Core Genotypes (FCG) founder male (XY^**-**^*Sry*^**+**^) harbors a Y chromosome with a deletion of the *Sry* gene, and a transgene encoding *Sry* on chromosome 3. Progeny of a cross between the XY^**-**^*Sry*^**+**^ male with a wild-type female results in offspring with four genotypes: two gonadal males (XY^**-**^*Sry*^**+**^ M, XX *Sry*^**+**^ M) and two gonadal females (XXF, XY^**-**^F). Hearts were obtained from 10.5 and 16.5 dpc embryos, neonates, and adults for analysis by RNA-seq and for CUT&RUN in the adult hearts. Differentially expressed genes (DEGs) and differential chromatin features (DCFs) influenced by sex chromosome composition and hormonal status were obtained for each stage.

To visualize global effects of sex-biasing factors on gene expression, we conducted principal component analysis (PCA) of the resulting cardiac transcriptomes. As expected, principal component analysis shows that developmental stage contributes most to the observed variance between samples (**Figure 3**).

**Figure 3.**
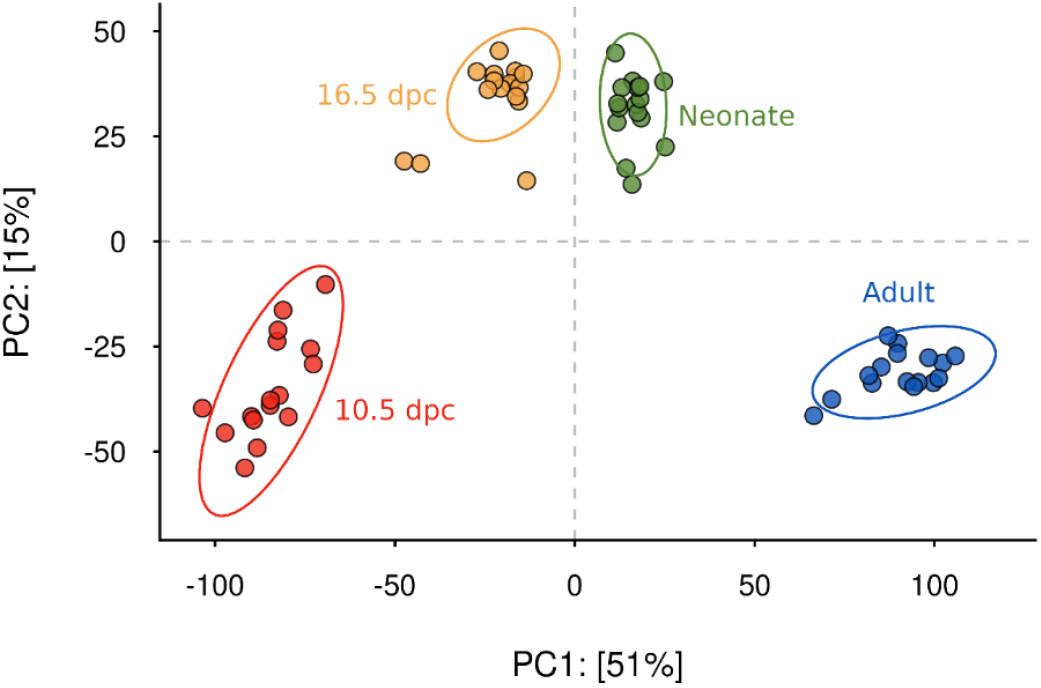
Principal component analysis of cardiac transcriptomes. PCA plot of the four cardiac stages analyzed shows that the major variable for cardiac gene expression for all four core genotypes is the developmental stage.

We validated our RNA-seq results by comparing our data with previously published transcriptional factors typical of early cardiac development^35-37^. For example, holding chromosomal sex and gonadal status constant, we found 1,971 differentially expressed genes between 10.5 and 16.5 dpc at FDR < 0.05. These included transcription factors more highly expressed at the 10.5 dpc stage such as *Isl1, Sall4, Eomes*, and *Gata1*, and more highly expressed at 16.5 dpc such as *Tcf15, Irf8*, and *Hoxd8*.

We tested for the effects of female *versus* male gonad types holding chromosomal sex and developmental stage constant and found 340 differentially expressed genes, including transcription factors *Esr1, Id2*, and *Klf9* higher in gonadal females and *Tcf15* and *Ybx2* higher in gonadal males. Of these, 90 genes exhibited a significant interaction effect, i.e., the male/female differential expression dependent on the sex chromosome status (**Supplemental Table 6**). Gene ontology terms for genes upregulated in females included glycosaminoglycan binding (molecular function), epithelium development (biological process), and extracellular matrix (cellular component), and the top upregulated pathway was epithelial-mesenchymal transition. Gene ontology terms for genes upregulated in males included vascular endothelial growth factor receptor 1 binding (molecular function), mitochondrial gene expression (biological process), and mitochondrial protein-containing complex (cellular component). Oxidative phosphorylation was the top upregulated pathway of genes more highly expressed in males, considering all developmental stages (**Supplemental Table 7**).

The sex chromosome effects holding gonad type and developmental stage constant yielded 174 differentially expressed genes, including *Hoxa4* and *Hoxa5*, higher in XX mice. *Tbx4l* exhibited differences with significant interaction depending on the gonadal status. Gene ontology terms for genes upregulated in XX mice included receptor ligand activity (molecular function), epithelium development (biological process), and extracellular matrix (cellular component), and the top upregulated pathway was xenobiotic metabolism. Gene ontology terms for genes upregulated in XY mice included dioxygenase activity (molecular function), supramolecular fiber organization (biological process), and I band (cellular component). Oxidative phosphorylation was the top upregulated pathway of genes more highly expressed in XY mice, considering all developmental stages.

### Sex chromosome and sex hormone effects on gene expression at each cardiac developmental stage

To distinguish between sex chromosome and sex hormone effects on sex-biased expression in the heart at each developmental stage, we computed the intersections of the contrasts due to presence or absence of *Sry* and those due to gonadal effects. We discuss genes that had an FDR <0.05 (**Figure 4**).

**Figure 4.**
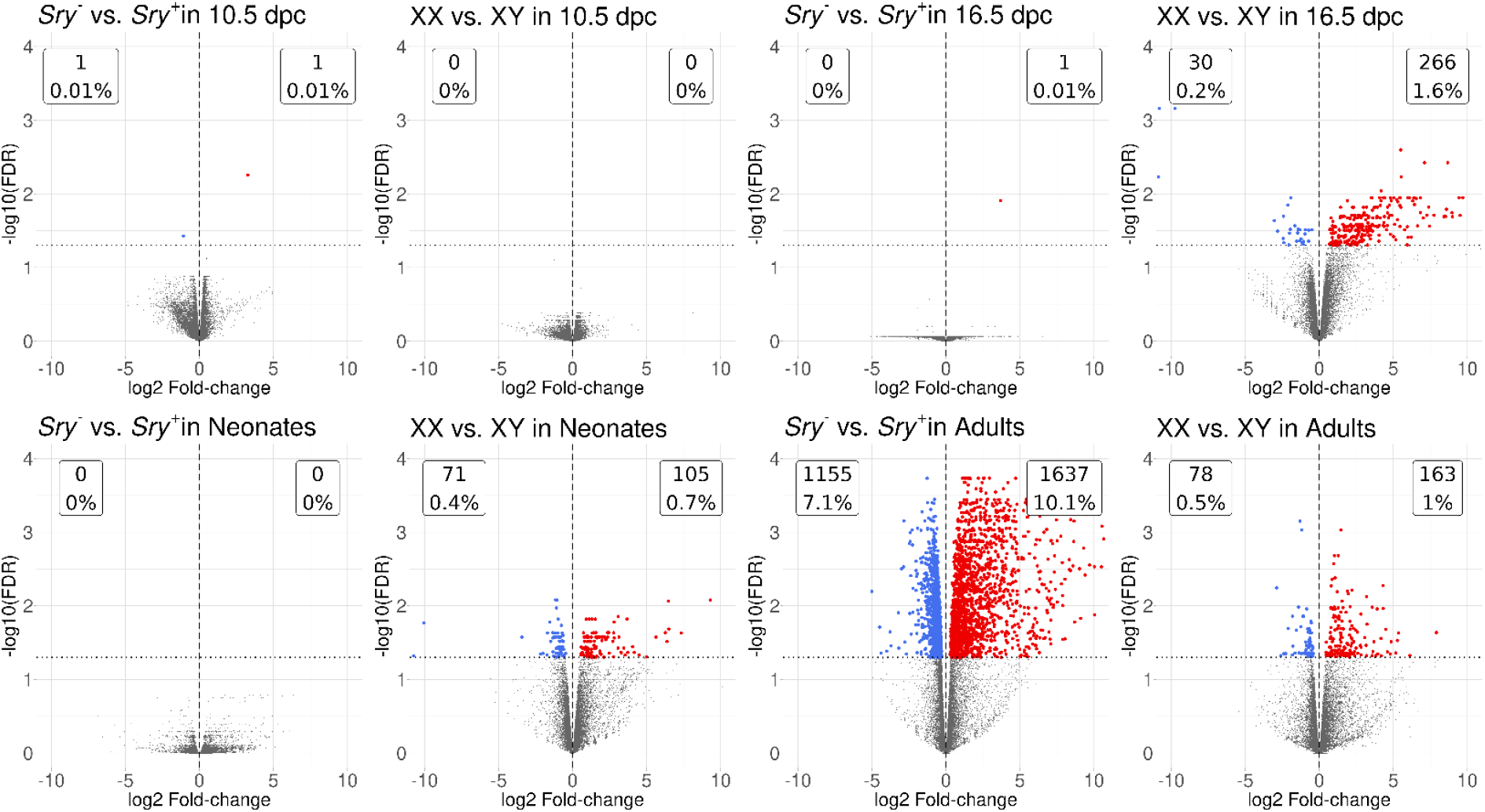
Analysis of sex chromosome and gonadal hormone effects for each cardiac developmental stage. Volcano plots showing the intersections of the contrasts due to presence or absence of *Sry* and those due to gonadal effects for hearts for 10.5 dpc and 16.5 dpc embryos, neonates, and adults.

At 10.5 dpc, sex-biased gene expression can be attributed solely to sex chromosome effects. However, the XX *versus* XY comparisons, whether for gonadal males or females, exhibited no significantly biased expression of either Y- or X-linked genes.

In 16.5 dpc embryos, 258 genes were differentially expressed depending on sex chromosome constitution, only 13 of which were linked to either the X or the Y chromosome. As expected, the long non-coding RNA *Xist* was observed to be expressed only in XX mice, whether gonadal males or females. Among the sex-biased genes, 22 transcription factors were more highly expressed in XX mice, including 6 members of the *Hox* family. Two Y-linked epigenetic factors, *Uty* and *Kdm5d*, were only expressed in XY mice, as expected. At this stage, none of the differentially expressed genes could be attributed to the independent contribution of gonadal hormones.

In neonates, 102 genes exhibited sex chromosome effects, 9 of which were either X- or Y-linked. None of the X-linked genes that have been reported to escape X chromosome inactivation^38^ were observed to be more highly expressed in XX gonadal females or males in the heart.

Interestingly, several X-linked genes are more highly expressed in XY mice, including *Alas2*, an encoding protein involved in porphyrin metabolism, and *Mid1*, an X-linked gene encoding a protein involved in anchoring other proteins to microtubules^39^. There were no significantly biased genes due to hormonal effects.

In adult hearts, 58 genes exhibited differential expression due to sex chromosome effects alone. Nine X-linked genes were more highly expressed in XX mice, including 3 genes reported to escape X chromosome inactivation (*Kdm5c, Ddx3x*, and *Eif2s3x*). *Kdm6a*, previously reported to escape X chromosome inactivation, was not more highly expressed in XX mice in the heart.

Furthermore, 2,563 genes were differentially expressed due exclusively to hormonal effects: 1,495, including *Esr1*, were more highly expressed in female mice and 1,068 in male mice. Ninety-one genes showed both sex chromosome and sex hormone contributions on their expression patterns. For example, *Wnt9b* was more highly expressed in XX *versus* XY mice and in gonadal females *versus* males. Both *Sox4* and *Irx1* were more highly expressed in XY *versus* XX mice and in gonadal males *versus* females.

### Sex-biased chromatin status

To identify regions of differential chromatin state between the adult hearts of FCG mice, we identified regions with differential enrichment of H3K27Ac marks, as these are associated with enhancer activity. The comparisons for differential abundance analysis of H3K27Ac marks were XX *versus* XY differences in gonadal females, XX *versus* XY differences in gonadal males, gonadal female *versus* gonadal male hormonal effects in XX genotype, gonadal female *versus* gonadal male hormonal effects in XY genotype, average difference in gonadal female *versus* gonadal male holding sex chromosomes constant, and average difference in XX *versus* XY holding hormones constant (**Figure 5**). The gonadal female *versus* gonadal male contrast showed 5 up-regulated (female-biased) and 84 down-regulated (male-biased) peaks, which indicate hormonal effects. The XX *versus* XY comparisons showed 195 up-regulated (XX-biased) and 90 down-regulated (XY-biased) peaks dependent on sex chromosome effects. (26 regions showed joint hormonal and sex chromosome effects).

**Figure 5.**
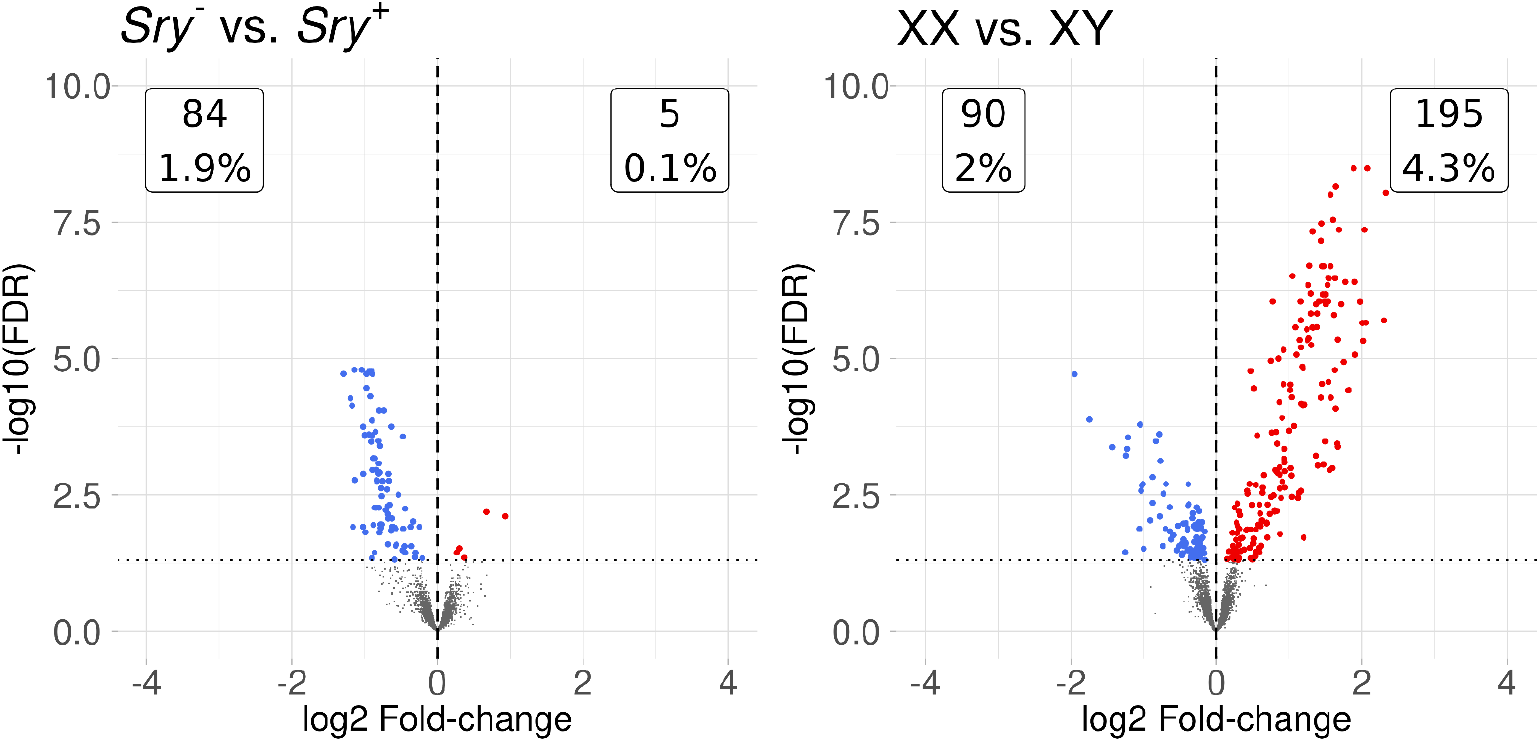
Volcano plots showing the results of differential abundance testing of the indicated contrasts for chromatin status. CUT&RUN was performed on adult hearts to identify regions of differential enrichment of H3K27Ac.

### Concordance between RNA-seq and CUT&RUN analysis

Differentially expressed genes from adult hearts were correlated with the nearest upstream CUT&RUN peak (**Figure 6**). Although we found no strong pattern of correlation for either the XX *versus* XY or male *versus* female comparisons, some genes were an exception (track plots shown in **Supplemental Figures 3-7**).

**Figure 6.**
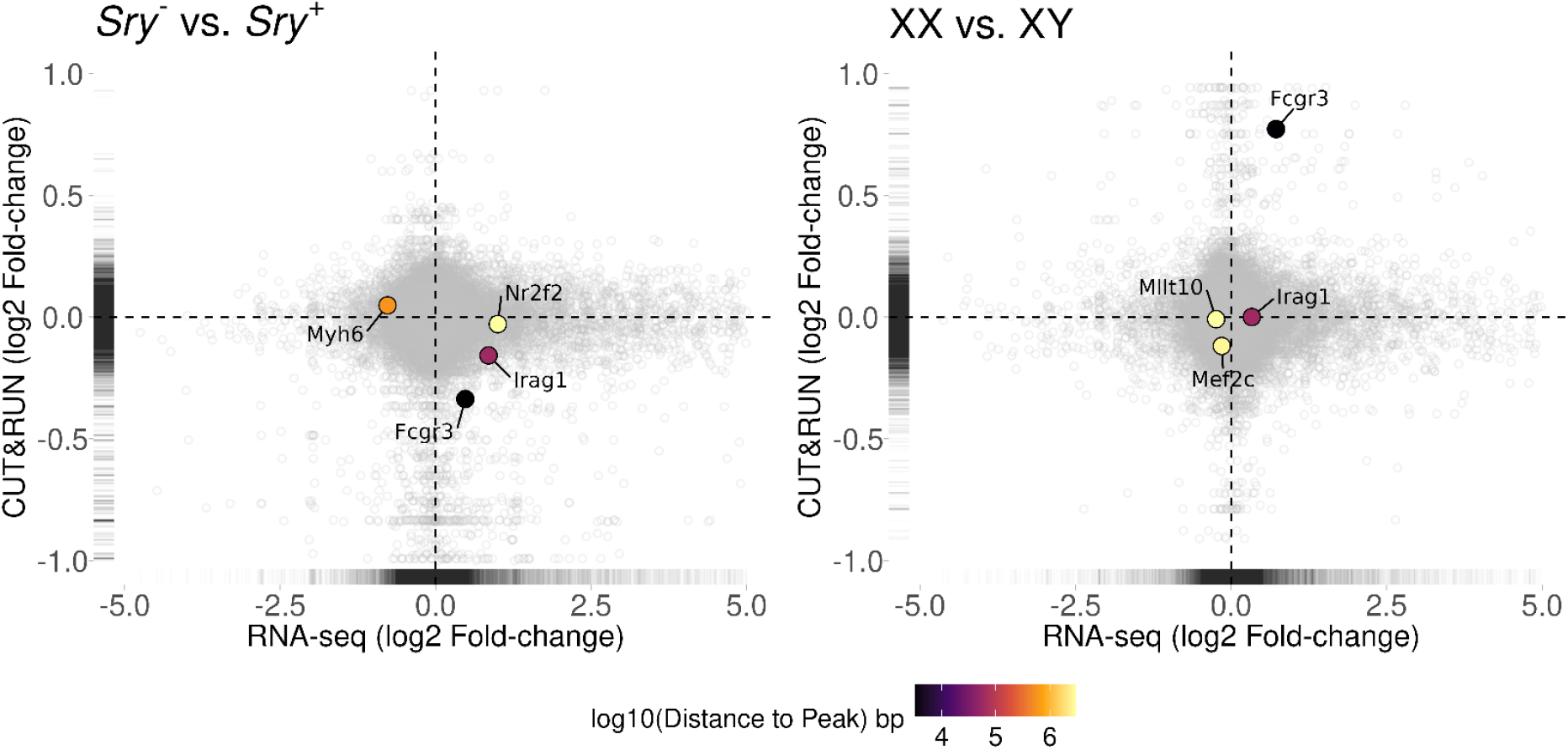
Concordance between RNA-seq and CUT&RUN analysis. The left and right panels show the logFC of gene expression between conditions (Left: female *versus* male; Right: XX *versus* XY) on the X axis and the logFC of the nearest upstream CUT&RUN peak on the Y axis. Genes relevant to cardiac development are highlighted.

For example, 4 genes had an expression change that correlated with a change in H3K27Ac abundance in a neighboring region that indicated a hormonal effect, 144 gene-peak correlations suggesting a sex chromosome effect, and 13 gene-peak correlations indicating a possible joint effect.

### Motifs present in enriched H3K27Ac regions

We performed motif analysis to determine whether the sequences with differentially accessible chromatin for each comparison were potential transcription factor binding sites in adult hearts of the FCG mice. The comparisons were made for regions with significantly higher enrichment for H3K27Ac for XX *versus* XY chromosome composition and males *versus* females. We found that regions with XX-biased H3K27Ac enrichment had significant similarity to *Err2, FoxJ3, Pitx1* and *Irf2*, among others. Regions with XY-biased open chromatin shared some of these motifs, but also included *HoxA1*. Female-biased H3K27Ac enriched regions had motifs significantly similar to *Atf3, Wt1*, and *Esr1/Esr2*. Conversely, regions with male-biased enrichment in H3K27Ac had motifs with similarity to *Ascl1*, but also *Esr1/Esr2*.

### CUT&RUN differential peaks overlapping known cardiac enhancers

We compared the sex chromosome- or sex hormone-dependent differential H3K27Ac peaks to regions containing known cardiac regulatory elements. We found 5 sex hormone-dependent and 80 sex chromosome-dependent regions overlapped with reported cardiac enhancer regions, with 5 differential H3K27Ac peaks appearing in both the F *versus* M and XX *versus* XY comparisons. Twenty-two genes associated with these overlapping regions exhibited cardiac phenotypes in mice.

## Discussion

Sex differences in gene expression occur in every tissue and result from differences in genotypic sex as well as sex differences in circulating and local hormonal regulators. Disentangling these contributing factors and identifying their independent effects, in addition to their interactions, has been achieved with genetically modified mice, such as the Four Core Genotypes (FCG) model, among others. The majority of these studies have used these models to elucidate how sex hormones and the sex chromosome composition affect diseases in adult mice^40-42^. By contrast, our goal was to leverage the FCG model to study a developmental process. We analyzed RNA expression patterns across cardiac development, beginning at 10.5 dpc and culminating in the adult. The sex biases in expression patterns that we observed lay the foundation for studying the developmental origins of cardiac phenotypes and diseases that exhibit sex differences.

Congenital heart disease is frequently sex-biased^1,2,43^, but these disparities have not been accounted for. The prevalence of congenital heart disorders in Turner Syndrome patients motivated the search for X-linked genes that could contribute to cardiac development. As one of the two X chromosomes in females undergoes inactivation, the focus has been on genes that escape silencing, are more highly expressed in females, and are dosage-dependent^44^. For example, having one copy of *TIMP1* (Xp11.3) in Turner Syndrome patients increases the risk of bicuspid aortic valve, coarctation of the aorta and aortopathy^45^. Haploinsufficiency of the escapee gene *KDM6A* (Xp11.3) causes congenital heart disease in Kabuki Syndrome^46^.

We used the FCG mouse model to determine when sex biases in transcriptional patterns appear during cardiogenesis and whether they are determined by sex chromosome or gonadal hormonal effects, or both. For all our data, we ignored the 9 genes included in the X-Y translocation that were differentially expressed. We find that cardiac cells have sex-biased RNA expression at 10.5 dpc, preceding gonad formation. These results confirm previously reported sex differences in RNA abundance and protein expression patterns at 9.5 dpc^47^ and underscore the importance of sex chromosome contributions to genome-wide sex-specific expression patterns in early developmental stages, in the absence of endogenous circulating sex hormones. Importantly, one Y-linked histone lysine demethylase, *Kdm5d*, expressed almost immediately post-fertilization^8^, could potentially establish genome-wide chromatin modifications that determine the availability of regulatory elements in subsequent developmental stages, including in the heart. While this hypothesis was proposed many years ago^12^, it remains to be tested experimentally.

The analysis of cardiac protein expression patterns at 9.5 dpc revealed sex-biased proteins that did not exhibit sex differences in RNA abundance, and some sex-biased RNAs did not result in sex-biased proteins^47^. Although not directly comparable to our 10.5 dpc data, this suggests that our list of sex-biased RNAs at this stage may underestimate the sex differences in protein abundance.

The vast majority of differentially expressed genes in the heart at 16.5 dpc are regulated by the sex chromosome composition, suggesting that although the gonads have differentiated and testes secrete more testosterone than ovaries by 13 dpc (PMID: 522476), hormonal effects are not yet evident. Surprisingly, 6 members of the homeobox gene family (*HoxA2, HoxA3, HoxA4, HoxA5, HoxB2, HoxB4*), and 3 members of the *Tbx* family (*Tbx1, Tbx2, Tbx4*) were more highly expressed in XX than XY mice, whether they were gonadal males or females. The ontology terms most significantly associated with XX-biased genes were related to morphogenesis and development, whereas for the XY-biased genes they were related to enzymatic processes and chromatin remodeling. This suggests that 16.5 dpc XX and XY hearts are at different developmental stages. Alternatively, because the data derives from whole hearts, these effects could reflect differences in cell type composition. The mechanism leading to this offset of cardiogenic stages is unknown, but clearly depends on the sex chromosome composition and could explain the sex differences in susceptibility to congenital heart disease. Previous studies have suggested that male embryos may be more advanced than females, at least in post-implantation embryos^48^. One possible explanation is that because of the need for X chromosome inactivation in XX mice, some developmental stages are slightly delayed in those mice. In mouse embryonic stem cells, XX cells lag behind XY and XO cells in their gene expression patterns upon differentiation to embryoid bodies^49^.

Surprisingly, we found that all differentially expressed genes in neonatal hearts occurred in the contrast between XX and XY mice. Ontology terms enriched in XX-biased genes were again related to developmental processes, but also negative regulation of proliferation, whereas for XY-biased genes they were associated with cell cycle and DNA replication.

We had expected to see hormonal effects on gene expression in neonatal hearts because males have a testosterone spike at that stage^52,53^. We hypothesize that, at least in the heart, there is insufficient expression of the androgen receptor to impose transcriptional differences dependent on the circulating androgens in gonadal male neonates. Another possible explanation is that the sex chromosome effects counteract or mask the hormonal effects.

The majority of sex-biased gene expression in the adult hearts was dependent on hormonal effects, as expected. These included genes associated with cardiac phenotypes, such as *Tbx5, Fgfr2, Smad6* and *Timp2*, among others.

Interestingly, genes for which expression differences were dependent on both hormonal and sex chromosome effects included transcription factors *Sox4* and *Irx1*, with higher expression levels in gonadal males and XY mice, whether male or female, and *Twist1* and *FoxP2* exhibiting higher levels in gonadal females and XX mice. The mechanisms by which these effects interact require further molecular studies.

We compared our data to the sex-stratified Gtex RNA-seq data from heart ventricle. The majority of sex-biased genes did not overlap between humans and mice. This is in keeping with data suggesting that, at least in somatic tissues, there is a fast evolutionary turnover of sex-biased expression^54^. Of the female-biased genes that overlapped between mice and humans, 2 had associated heart phenotypes, *Plagl1* and *Sema3c*. None of the male-biased genes exhibited abnormal cardiac phenotypes in the mouse. Nevertheless, because of strong conservation of X and Y genes that are implicated in causing sex differences in mice, our studies suggest that sex chromosome genes are very likely agents causing sex differences during early stages of cardiac development in numerous mammalian species, including humans.

Taken together with a previously published report^47^, our data show that sex differences in gene expression are apparent in cardiac development well before gonad formation, and thus, established by the differences in sex chromosome composition. Because many congenital heart diseases are sex-biased, further exploration will be necessary to associate those disorders with sex-specific networks.

Expression differences between the sexes continue to exist in the heart throughout the lifespan, with gonadal hormones playing a major role in regulating these sex biases, independently and also in jointly with the sex chromosomes. Our data does not uncover interactions in which the two factors exert opposing effects, canceling each other out. However, our experiments provide a valuable resource that yields potential candidates, including specific genes and regulatory regions, for studying the sex biases that occur in the incidence and presentation of adult cardiac diseases and a model for studying the sex disparities observed in many other diseases.

## Acknowledgements

We thank Kelsey Keith, Kayla Johnson, and Arjun Krishnan for their analyses and input.

---

**Supplemental Table 1:**
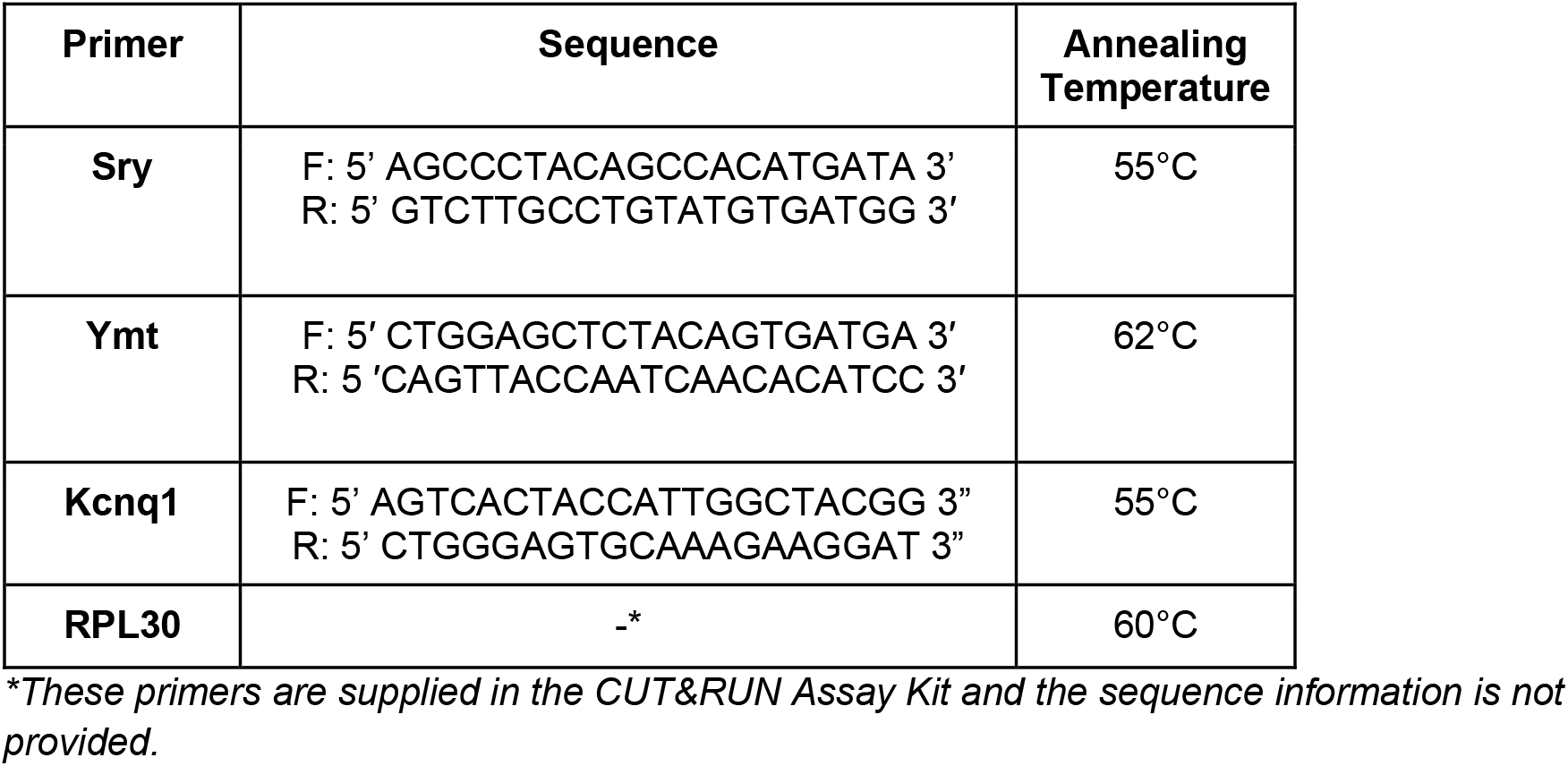
Primer Sequences.

## Supplemental Figures

**Supplemental Figure 1.**
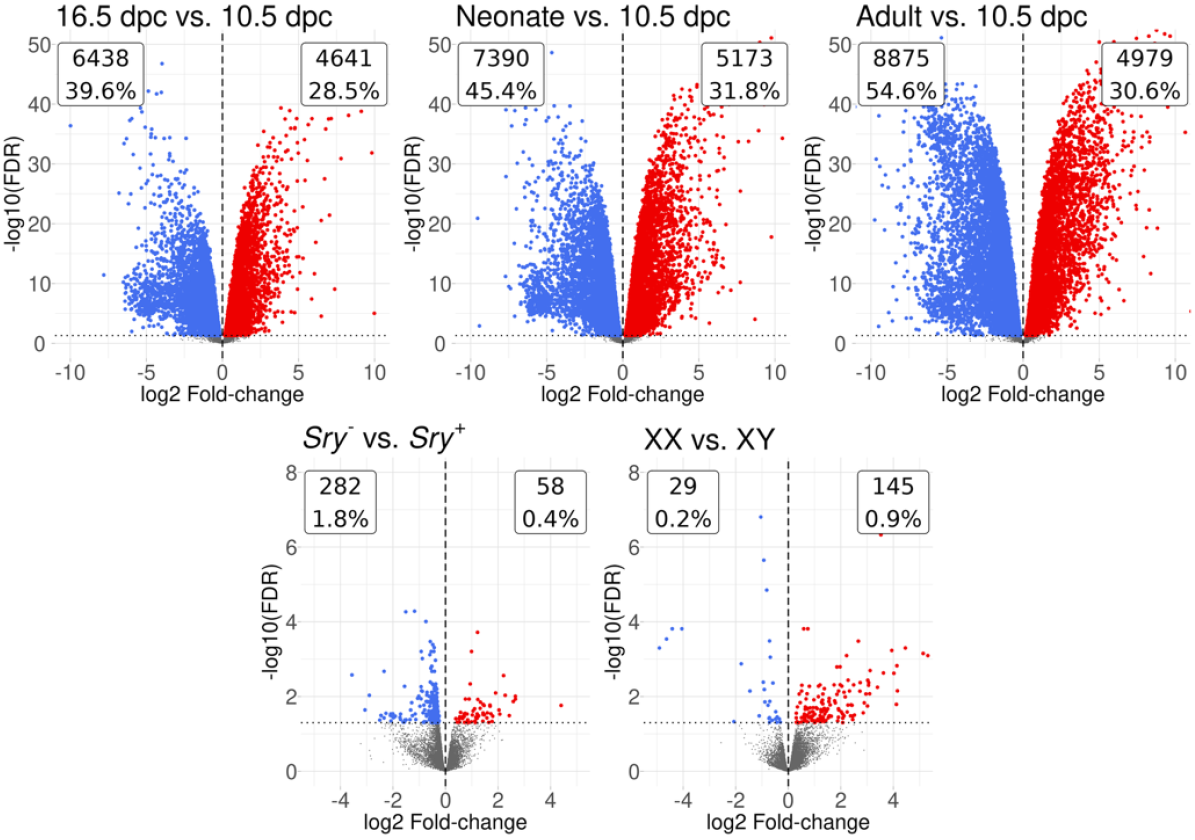
Volcano plots showing the intersections of the contrasts due to presence or absence of *Sry* and those due to gonadal effects for hearts for 16.5 dpc embryos, neonates, and adults compared to 10.5 dpc hearts.

**Supplemental Figure 2.**
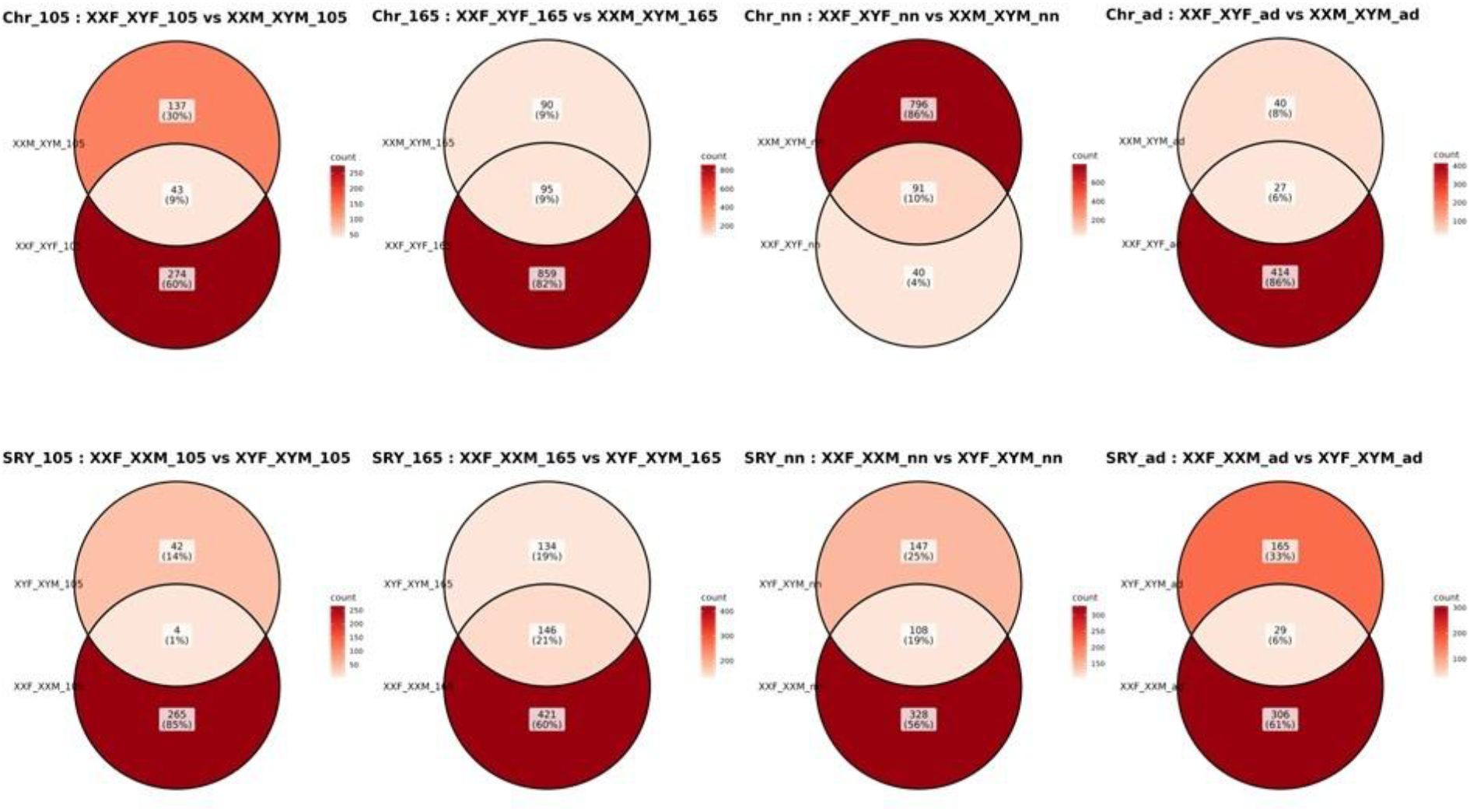
Intersection analysis of gene numbers for sex chromosome (upper panel) and gonadal status (lower panel) for each cardiac stage.

**Supplemental Figure 3.**
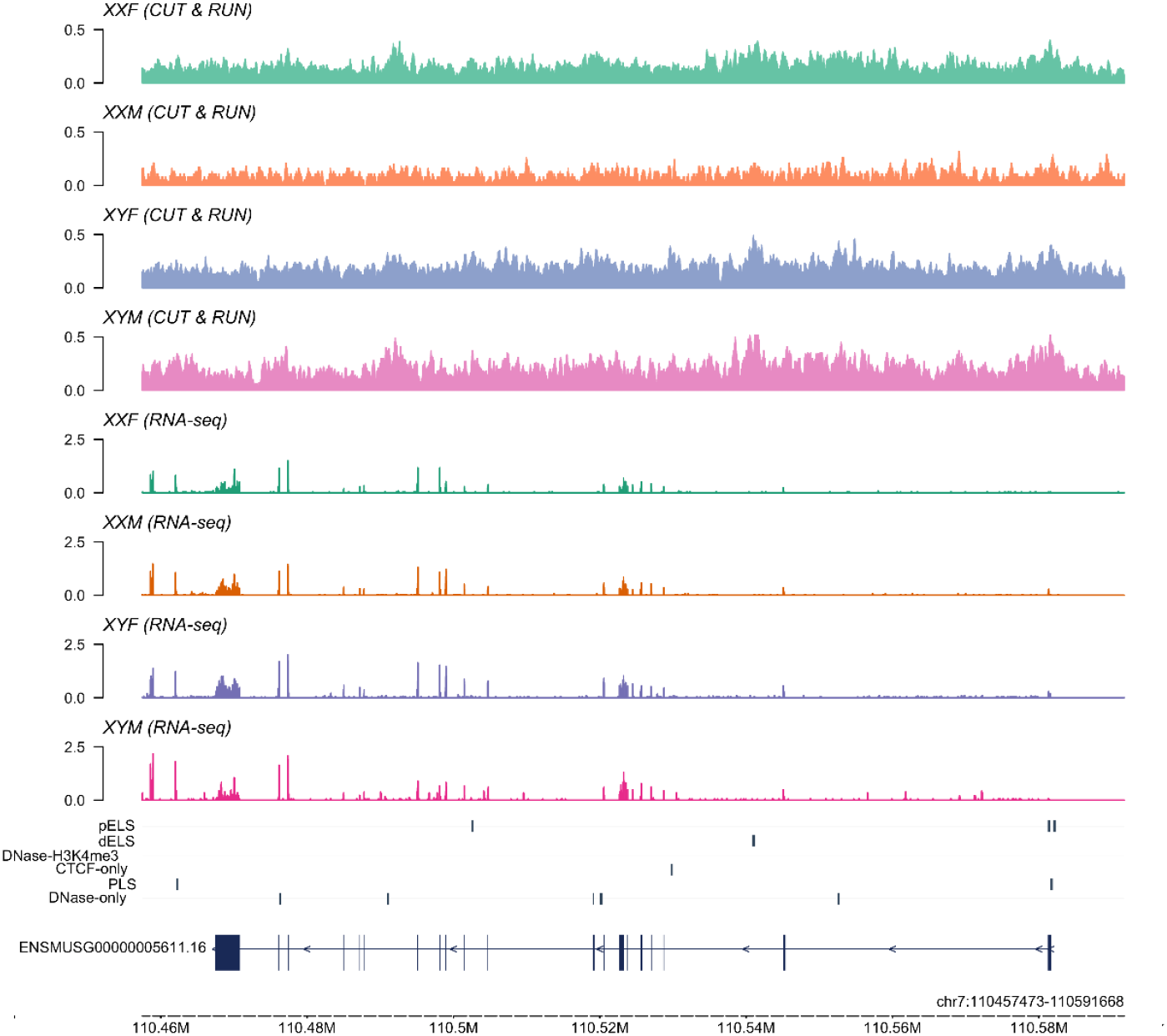
Track plots of H3K27Ac peaks and RNA-seq results for *Irag1*.

**Supplemental Figure 4.**
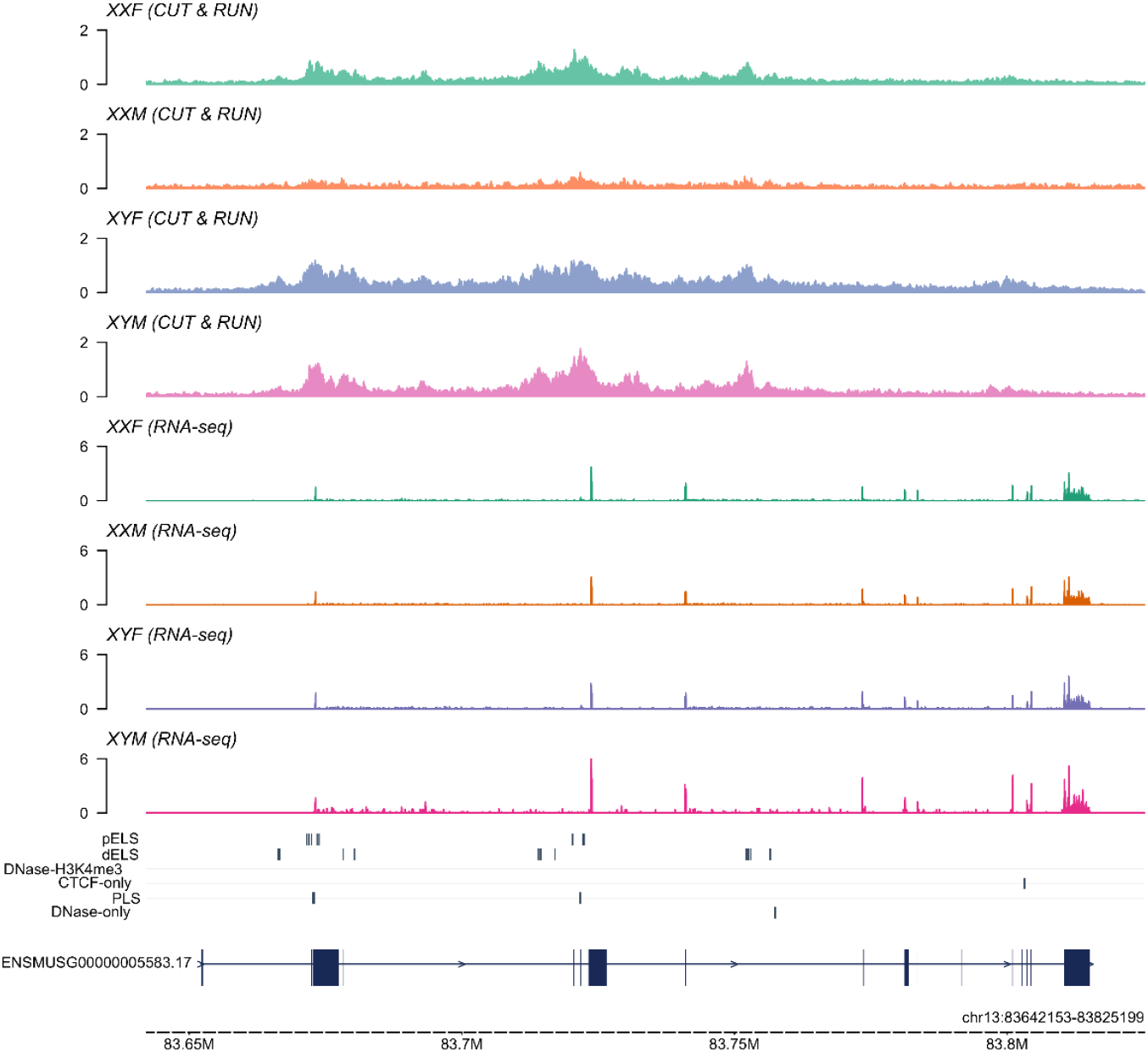
Track plots of H3K27Ac peaks and RNA-seq results for *Mef2c*.

**Supplemental Figure 5.**
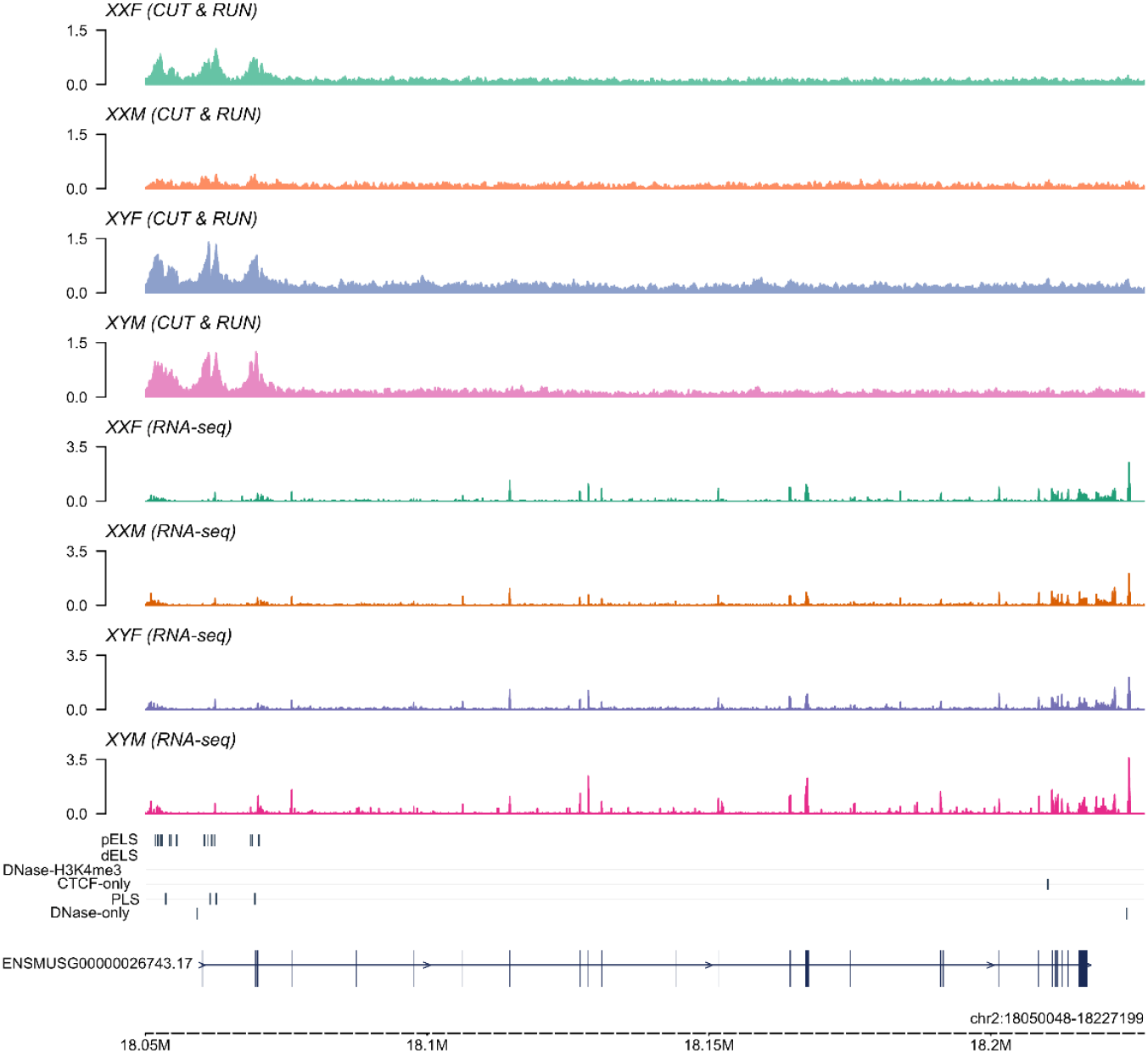
Track plots of H3K27Ac peaks and RNA-seq results for *Mllt10*.

**Supplemental Figure 6.**
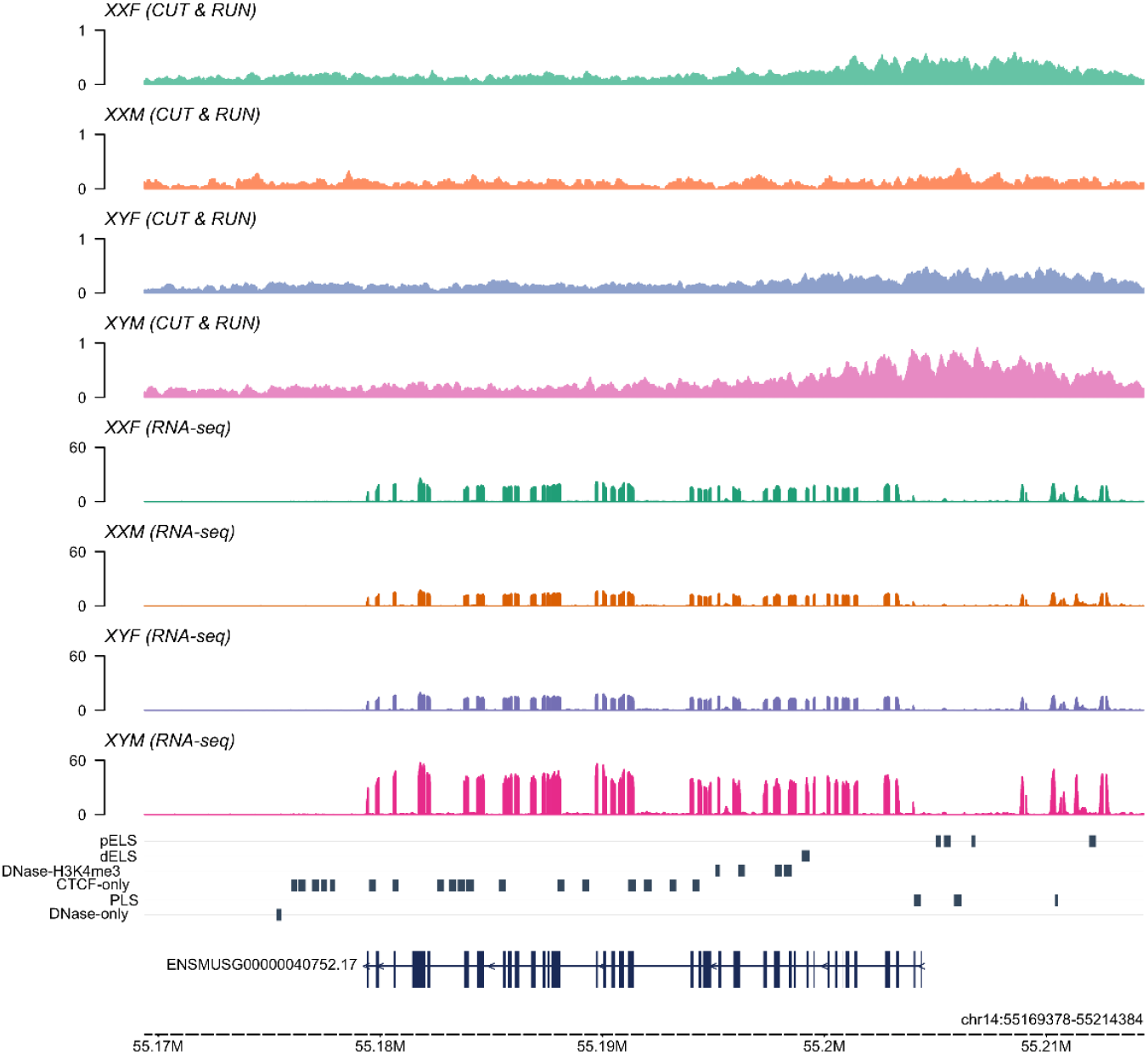
Track plots of H3K27Ac peaks and RNA-seq results for *Myh6*.

**Supplemental Figure 7.**
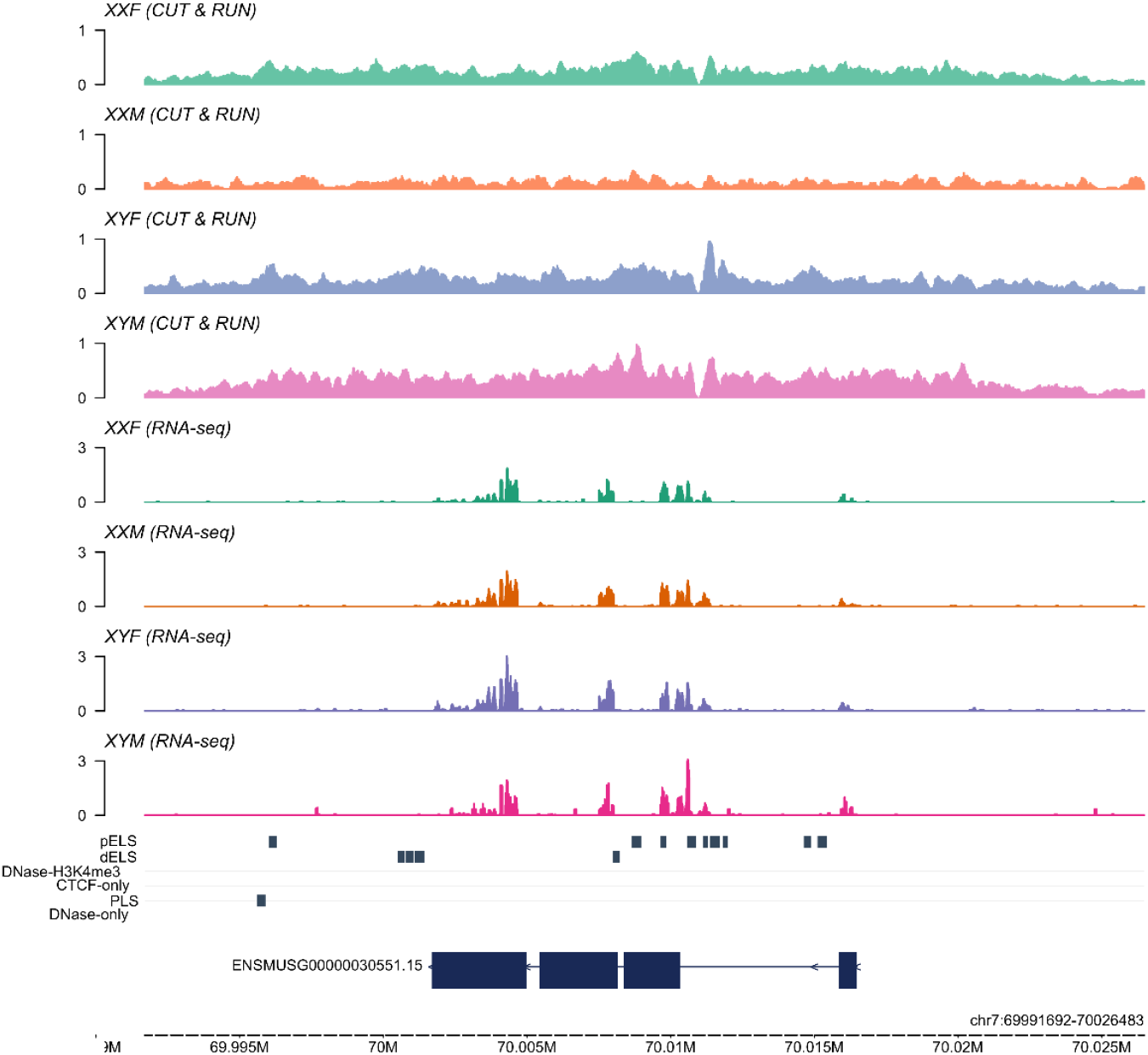
Track plots of H3K27Ac peaks and RNA-seq results for *Nr2f2*.

